# Familiarity and Face-Inversion Effect in Japanese Macaques (*Macaca fuscata*) during the Preferential Looking Task

**DOI:** 10.1101/267716

**Authors:** Masaki Tomonaga

## Abstract

Four young laboratory-born Japanese macaques (*Macaca fuscata*) looked at the photographs of familiar and unfamiliar persons presented at upright and inverted orientations by pressing the lever under the conjugate schedule of sensory reinforcement (successive preferential looking procedure). Three types of photographs were prepared: photographs with persons taken in front view, those taken in back, and those without persons. The monkeys looked longer when the face was upright than inverted only for the pictures containing unfamiliar person with front view. The other types of photographs did not cause inversion effect. Familiarity weakened the face-specific inversion effect in monkeys. This difference may be due to in part the lower preference for familiar faces and the difference in processing mode between familiar and unfamiliar faces.

## INTRODUCTION

In humans perception of inverted faces is severely impaired when the faces are presented at inverted orientation (Bruce, 1988; Yin, 1969). This inversion effect is due to the difference in processing between upright and inverted faces: configural properties are utilized for the processing of upright faces, while individual features for inverted faces (Carey & Diamond, 1977; Diamond & Carey, 1986). Recent advances accumulated the evidence *for* face-inversion effect in nonhuman primates (Phelps & Roberts, 1994; Wright & Roberts, 1996; Parr, Dove, & Hopkins, 1998; Tomonaga, 1994, 1999, 2018). For example, Tomonaga (1994) examined face inversion effect in laboratory-raised Japanese macaques (*Macaca fuscata*) under the modified preferential looking task. Monkeys were allowed to look at the photographs during they held down the lever. They showed significant difference in preference assessed with looking time between upright rhesus and Japanese macaque photographs, whereas no difference when photographs were presented at 90 or 180 degrees. This inversion effect was the strongest for the photographs containing front faces clearly. When photographs contained no clear faces, the inversion effect disappeared. Additional experiments with scrambled photographs in which inverted face with upright body or vice versa appeared also supported the Tomonaga’s results. Inversion effect was governed by the orientation of face but not by the orientation of body and background (Tomonaga, 2018).

There are some other effects on face perception in humans (Bruce, 1988) such as other-race effect (Malpass & Kravitz, 1969) and distinctiveness effect (Valentine, 1991). Furthermore, it is well known that familiarity also affects the face perception. Humans showed better recognition accuracy for familiar faces than for unfamiliar faces (Bruce, 1988; Young, McWeeny, Hay, & Ellis, 1986). It might be plausible that the familiarity also affect the inversion effect. Actually, familiar faces were little affected by the inversion during the naming task in the chimpanzee (Tomonaga, Itakura, & Matsuzawa, 1993). In the present experiment, the Japanese macaques were presented the photographs of familiar and unfamiliar humans during the successive preferential looking task to investigate the role of familiarity in the inversion effect.

## METHODS

### Participants

Four young Japanese macaques participated in the present experiment. At the onset of the present experiment, they were 5 years old in average (range: 4 to 6). They had been isolated from their mothers within one week after birth and raised by human caretakers. They lived in a cage (70 × 70 × 70 cm) with another macaque. They also participated in various types of sensory-reinforcement experiments (Fujita, 1990; Tomonaga, 1994, 2018). During the present study, they were not deprived of either food or water. This study was approved by the Animal Welfare and Care Committee of the Primate Research Institute, Kyoto University, and followed Guidelines for the Care and Use of Laboratory Primates of the Primate Research Institute, Kyoto University.

### Apparatus

The present experiment was conducted in an experimental room, in which the two identical chambers (60 × 60 x 60 cm) were set up. The front panel (40 × 40 cm) of each chamber was made of clear Plexiglass. A slide projector (CABIN, model Family Cabin) was installed behind the screen and projected the slide photographs onto a 33 × 33 cm opaque screen placed 50 cm apart from the front panel. A single lever with a red lamp was installed in the lower center of the front panel. All experimental events were controlled and recorded by MSX2 personal computers (TOSHIBA, model HX-34).

### Stimuli

Three types of color photographs of humans were prepared. Figure 1 shows examples of each type of photographs. The first type contained a single human person taken in a front view with various backgrounds (FRONT). The second contained a person standing in the back (BACK). The third type of photographs contained only the background (NO PERSON). The last two types were used as control photographs to verify that inversion effect only occurred in photographs with front faces. Each type of photographs was divided into the two subgroups based on familiarity. Each type of photographs consisted of 30 different persons, 10 of which were familiar to the participants such as the experimenter (see Figure 1), former experimenters, caretakers, and the stuff around the experimental room who saw these monkeys very often. The other 20 persons were completely unfamiliar to the participants. BACK and NO PERSON photographs were also classified on the basis of this distinction. These photographs were presented at the two orientations, upright and inverted. Furthermore, two types of control slides (10 no light and 10 white light slides) were also presented, so that, 200 slides in total were prepared (180 photographs and 20 control slides). These 200 slides were randomly ordered and set in the two 100-slot circular cartridges.

**Figure 1.**
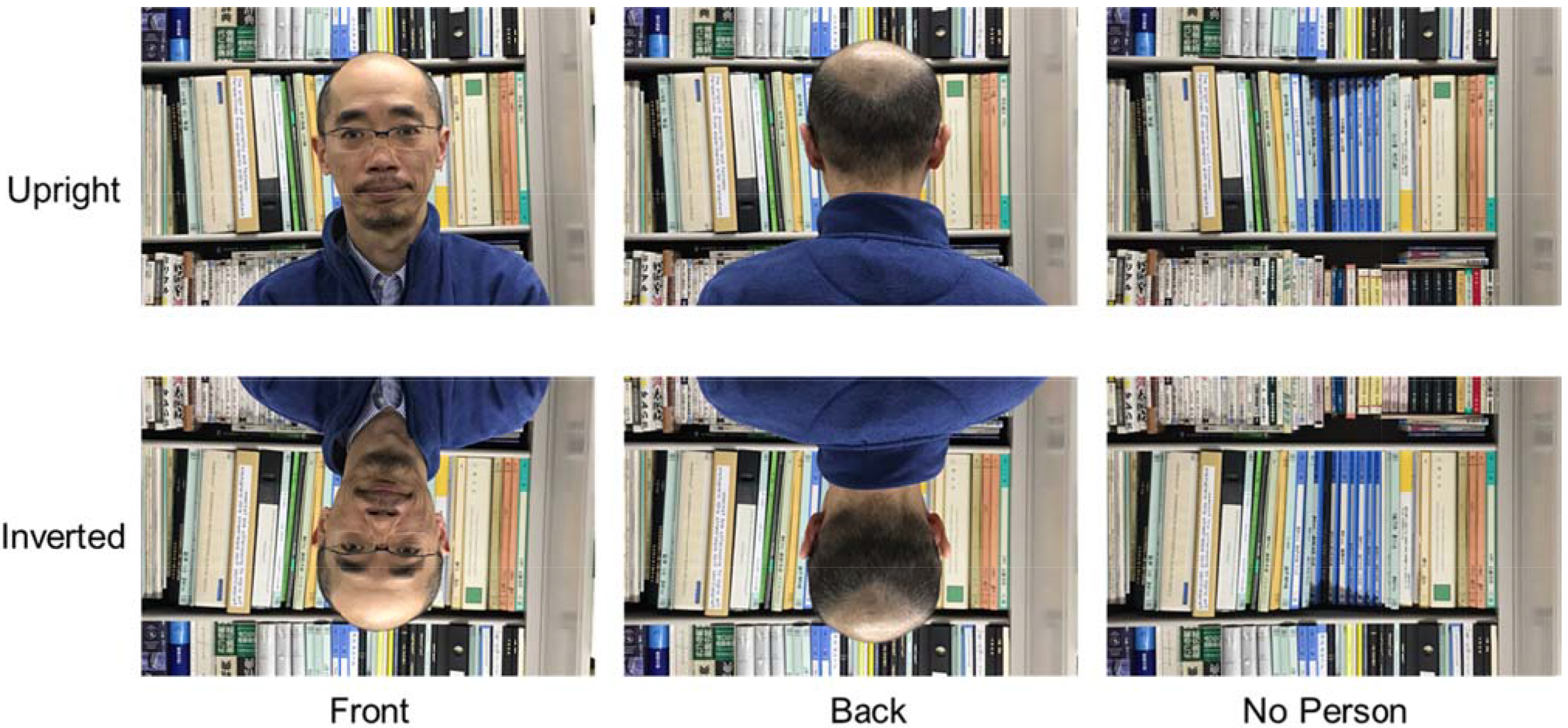
Examples of photographs of upright and inverted familiar person (the author).

### Procedure

In the present experiments, the modified version of preferential looking task was employed (Fujita, 1990; Tomonaga, 1994, 2018). When the monkey held down the lever in the presence of red light, the slide was projected onto the screen during the monkey’s holding of the lever. If the monkey released the lever or the lever holding exceeded 10 s, the slide was terminated. Very short response duration (faster than 100 ms) had no effect. When the monkey held down the lever again within the 10 s from the last lever release, the same slide was presented again. On the other hand, if the monkey did not press the lever more than 10 s, the next slide was set up for presentation. This interval was called interresponse interval. The trial was defined as the presentations of the same slide. Since the slide presentation was contingent upon the monkey’s response, it was considered as sensory reinforcement (Fujita & Matsuzawa, 1986). Since the duration of the trial was dependent upon the monkey’s responses, this task was also considered as “successive” preferential looking task. The strength of sensory reinforcement, or preference index for each slide was identified by measuring the response duration (D) and interresponse interval (I). The index was calculated with the formula below.

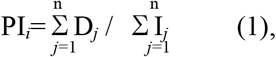

where PI_*i*_ designated the preference index of the *i*-th trial, n designated the number of responses in the *i*-th trial, D*j* designated the response duration of the *j*-th response, and I*j* designated the interresponse interval after the *j*-th response. When this index was higher, the monkey looked at the photograph for longer time and looked again with shorter interval. Because of the between-participants variances of the preference indices, the normalized index for each participant calculated with the formula (2) was used for the data analyses.

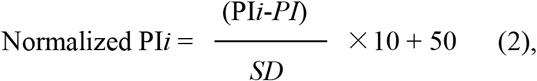

where the PI*i* designated the preference index for the slide *i*, *PI* and *SD* designated the mean and standard deviation of the preference indices for all slides. By normalizing, the mean value of preference index for each monkey was equated to 50.

Each slide was presented five times. The preference index for each slide was averaged across these five trials. These indices were normalized and then averaged for each type of photographs. A session was continued until the monkey completed 500 trials, that is five rotations of one cartridge. Each monkey received for two sessions (i.e. one session for one cartridge). The duration of each session was approximately 6 hours.

## RESULTS

Figure 2 shows the normalized preference index for each condition averaged across monkeys. For the difference in the preference between unfamiliar and familiar faces, 3 out of 4 monkeys showed higher index for upright unfamiliar than familiar FRONT photographs [55.51 vs. 52.26; randomization test (Edgington, 1987), *p*=0.0625 (1/16)].

**Figure 2.**
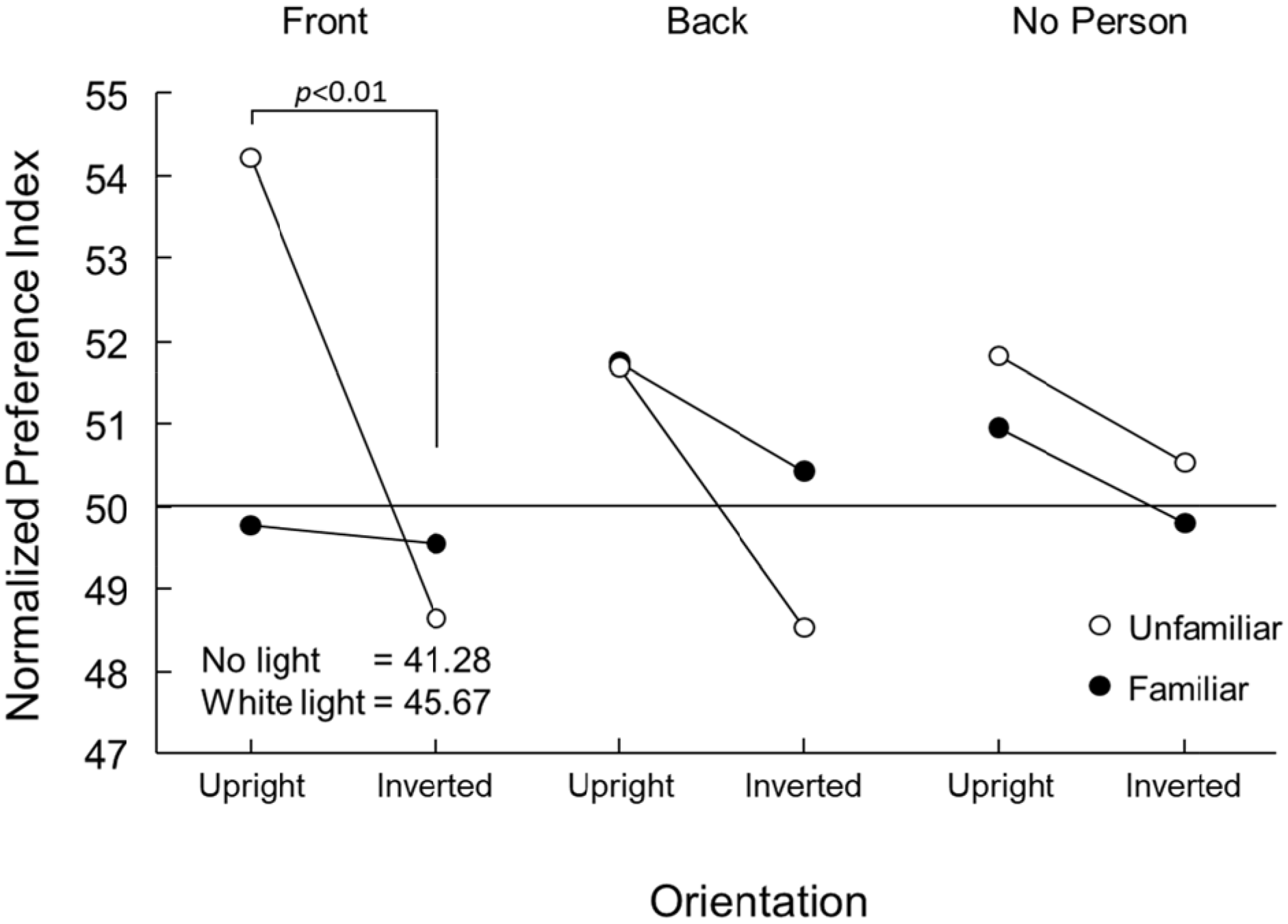
Normalized preference index for each type of photographs averaged across subjects. Values for the two control slides are shown as dotted lines.

Three-way analysis of variance (ANOVA) with repeated measures was conducted to the normalized preference indices. Main effects of photograph type and familiarity were not significant (*F*s<1), while the effects of orientation was significant [*F*(1,3)=27.80, *p*<0.05]. There were no significant two-factor interactions (*F*s<1.38), while triple interaction was significant [*F*(2,6)=9.43, *p*<0.05]. Tukey’s HSD tests revealed that lower preference index for inverted photographs than upright was only significant for unfamiliar FRONT pictures (*p*<0.01). Inversion effect was not significant for the other conditions.

Monkeys looked at the FRONT pictures of familiar persons (averaged across orientations) longer than the simple white light [randomization test *p*=0.0625 (1/16)]. This difference suggests that the loss of inversion effect for familiar persons was not derived from the overall floor effect.

## DISCUSSION

In the present experiment, the monkeys showed significant effect of inversion only when the photographs contained faces of unfamiliar humans. When the persons were present but no faces appeared (BACK photographs), and when only the background was presented (NO PERSON photographs), this effect was not evident. These results were consistent with the previous experiments by Tomonaga (1994, 2018). Inversion effect assessed with the preferential looking was specific to facial stimuli.

More important results of the present experiment are the difference in inversion effect between familiar and unfamiliar faces. The monkeys showed no inversion effect for familiar faces. In the present experimental condition, small difference in preference index between photographs indicates the small difference in strength of sensory reinforcement. Obviously, upright faces have reinforcing properties for monkeys (Fujita & Matsuzawa, 1986; Tomonaga, 1994; see also Kaplan, Fox, & Huckeby, 1992). Not so strong, but the present participants showed higher preference for upright unfamiliar faces than familiar faces. And the level of the index for familiar faces was comparable to those of control photographs. Familiar faces had less strong reinforcing properties than unfamiliar faces. The low preference to these photographs might have masked the inversion effect. This was not due to the overall floor effect, but relative one because the index for familiar faces was higher than simple white light but as low as control photographs. The lower preference for familiar faces is rather puzzling in comparison with the previous studies with human infants. In the sensory reinforcement experiments using sucking responses, human infants often showed consistent preference for their mother’s faces (most familiar to the babies) than strangers (Walton, Bower, & Bower, 1992). The dissociation between the present results and human infant studies might come from the qualitative difference of familiarity between mother and experimenters. The other possibility is that the preference index in the present experiment might reflect the other responses by the monkeys in addition to preference. It is often observed in everyday laboratory life that the monkeys more strongly and aggressively react to the strangers than familiar caretakers. The monkeys might have become alert when upright unfamiliar faces were presented.

In spite of the asymmetry of preference for familiar and unfamiliar faces observed in monkeys, possibly they could discriminate the familiarity among faces. As noted in Introduction, it is very easy for humans to judge the familiarity of faces (Bruce, 1988; Young et al., 1986). Chimpanzees can also recognize familiarity (Boysen & Berntson, 1986). From the attachment study, it is obvious that infant primates distinguish familiar and unfamiliar individuals (Inoue, Hikami, & Matsuzawa, 1993; Iue, Tomonaga, Zanma, & Kawakami, 2000). Then, what properties did the monkeys utilized for judgement of familiarity? And were those properties affected by inversion? As noted earlier, upright and inverted faces might be processed in different ways. However, the difference in processing mode between familiar and unfamiliar faces is still controversial (Carey & Diamond, 1994; Parr et al., 1998). Tomonaga et al. (1993) reported that the chimpanzee did not show the inversion effect during the naming task of familiar individuals. If this result is only affected by the familiarity (Parr et al., 1998; see also Tomonaga, 1999), it might be possible that their chimpanzee recognized familiar faces on the basis of differences in features which were robust to inversion. Usually, unfamiliar faces are dominantly processed in right hemisphere (e.g., Sergeant, 1988; Yin, 1970). However, some reports provided evidence that processing of famous faces showed left-hemispheric advantage (Marzi & Berlucchi, 1977). During the task in which the difference among faces was defined by the difference in facial parts, humans showed left-hemispheric advantage (Ross-Kossak & Turkewitz, 1984). Inui and Miyamoto (1984) found that in spite of including only local features, a familiar facial photograph could be perfectly identified by human observers. These results indicated that familiar faces were processed based on unique features (see also Parr et al., 1998). On the other hand, recent studies proposed both familiar and unfamiliar faces were processed configurally (Ellis, Shephard, & Davies, 1979; Young, Hellawell, & Hay, 1987; Carey & Diamond, 1994; Valentine, 1991).

If the present results suggested the loss of inversion effect in familiar faces, the monkeys might have processed familiar and unfamiliar faces in different ways: configural processing for unfamiliar (causing the inversion effect) and piecemeal or featural processing for familiar faces (robust for inversion). However, as suggested above, it is possible that familiarity itself lowered the looking time, so that, masked the inversion effect. It might be possible these two factors interactively affected the reduced inversion effect for familiar faces.

The present results are inconclusive about the processing mode of familiar faces. To investigate further the difference in processing mode between familiar and unfamiliar faces, future research should be conducted under the other discrimination tasks such as familiarity decision (Valentine, 1991), simple discriminations (Phelps & Roberts, 1994), or matching to sample (Parr et al., 1998; Tomonaga, 1999).

## ACKNOWLEDGMENTS

This study was financially supported by the JSPS/MEXT KAKENHI (#4710053, #13610086, #20002001, #23220006, #24000001, #15H05709, #16H06283), JSPS-CCSN and the JSPS Leading Graduate Program in Primatology and Wildlife Science (U04) at Kyoto University. The author wishes to thank Drs. Kazuo Fujita and Tetsuro Matsuzawa, and the staff of the Primate Research Institute, Kyoto University for their invaluable comments on this study.

## REFERENCES

Boysen, S. T.; Berntson, G. G. Cardiac correlates of individual recognition in the chimpanzee (Pan troglodytes). JOURNAL OF COMPARATIVE PSYCHOLOGY, 100: 321–324, 1986.

Bruce, V. RECOGNISING FACES. Hillsdale, NJ, Erlbaum, 1988.

Carey, S.; Diamond, R. From piecemeal to configurational representation of faces. SCIENCE 195: 312–314, 1977.

Carey, S.; Diamond, R. Are faces perceived as configurations more by adults than by children? VISUAL COGNITION 1: 253–274, 1994.

Diamond, R.; Carey, S. Why faces are and are not special: An effect of expertise. JOURNAL OF EXPERIMENTAL PSYCHOLOGY: GENERAL 115: 107–117, 1986.

Edgington, E. S. RANDOMIZATION TESTS (2nd ed.). New York, Marcel Dekker, 1987.

Ellis, H. D.; Shepherd, J. W.; Davies, G. M. Identification of familiar and unfamiliar faces from internal and external features: Some implications for theories of face recognition. PERCEPTION 8: 431–439, 1979.

Fujita, K. Species recognition by five macaque monkeys. PRIMATES 28: 353–366, 1987.

Fujita, K. Species preference by infant macaques with controlled social experience. INTERNATIONAL JOURNAL OF PRIMATOLOGY 11: 553–573, 1990.

Fujita, K.; Matsuzawa, T. A new procedure to study the perceptual world of animals with sensory reinforcement: Recognition of humans by a chimpanzee. PRIMATES 27: 283–291, 1986.

Inoue, N.; Hikami, K.; Matsuzawa, T. Attachment behavior and heart-rate change of infant chimpanzee (Pan troglodytes). JAPANESE JOURNAL OF DEVELOPMENTAL PSYCHOLOGY, 3: 17–24, 1992. (in Japanese with English summary)

Inui, T.; Miyamoto, K. The effect of changes in visible area on facial recognition. PERCEPTION 13: 49–56, 1984.

Iue, A.; Tomonaga, M.; Zanma, A.; Kawakami, K. Development of attachment between nursery-reared Japanese macaque infants and their human caregivers. PRIMATE RESEARCH 16: 274, 2000. (Japanese summary only)

Kaplan, P. S.; Fox, K. B.; Huckeby, E. R. Faces as reinforcers: Effects of pairing condition and facial expression. DEVELOPMENTAL PSYCHOBIOLOGY 25: 299–312, 1992

Malpass R. S.; Kravitz J. Recognition for faces of own and other race. JOURNAL OF PERSONALITY AND SOCIAL PSYCHOLOGY 13: 330–334, 1969.

Marzi, C. A.; Berlucchi, G. Right visual field superiority for accuracy of recognition of famous faces in normals. NEUROPSYCHOLOGIA 15: 751–756, 1977.

Parr, L. A.; Dove, T.; Hopkins, W. D. Why faces may be special: Evidence of the inversion effect in chimpanzees. JOURNAL OF COGNITIVE NEUROSCIENCE 10: 615–622, 1998.

Phelps, M. T.; Roberts, W. A. Memory for pictures of upright and inverted primate faces in humans (Homo sapiens), squirrel monkeys (Saimiri sciureus), and pigeons (Columba livia). JOURNAL OF COMPARATIVE PSYCHOLOGY 108: 114–125, 1994.

Ross-Kossak, P.; Turkewitz, G. Relationship between changes in hemispheric advantage during familiarization to faces and proficiency in facial recognition. NEUROPSYCHOLOGIA 22: 471–477, 1984.

Sergeant, J. Face perception and the right hemisphere, pp. 108–131 in THOUGHT WITHOUT LANGUAGE. Weiskrantz, L. ed. Oxford, UK, Clarendon Press, 1988.

Tomonaga, M. How laboratory-raised Japanese monkeys (*Macaca fuscata*) perceive rotated photographs of monkeys: Evidence for an inversion effect in face perception. PRIMATES 35: 155–165, 1994.

Tomonaga, M. Inversion effect on perception of human faces in a chimpanzee (Pan troglodytes). PRIMATES 40: 417–438, 1999.

Tomonaga, M. Inverted face with upright body: Further evidence for face inversion effect in Japanese macaques (Macaca fuscata). BIORXIV, 266676. 10.1101/266676.

Tomonaga, M.; Itakura, S.; Matsuzawa, T. Superiority of conspecific faces and reduced inversion effect in face perception by a chimpanzee. FOLIA PRIMATOLOGICA 61: 110–114, 1993.

Valentine, T. A unified account of the effects of distinctiveness, inversion, and race in face recognition. QUARTERLY JOURNAL OF EXPERIMENTAL PSYCHOLOGY 43A: 161–204, 1991

Wright, A. A.; Roberts, W. A. Monkey and human face perception: Inversion effects for human faces but not for monkey faces or scenes. JOURNAL OF COGNITIVE NEUROSCIENCE 8: 278–290, 1996.

Walton, G. E.; Bower, N. J.; Bower, T. G. Recognition of familiar faces by newborns. INFANT BEHAVIOR AND DEVELOPMENT 15: 265–269, 1992.

Yin, R. K. Looking at upside-down faces. JOURNAL OF EXPERIMENTAL PSYCHOLOGY 81: 141–145, 1969.

Yin, R. K. Face recognition by brain-injured patients: A dissociable ability? NEUROPSYCHOLOGIA 8: 395–402, 1970.

Young, A. W.; McWeeny, K. H.; Hay, D. C.; Ellis, A. W. Matching familiar and unfamiliar faces on identity and expression. PSYCHOLOGICAL RESEARCH 48: 63–68, 1986.

